# Chimeric synthetic reference standards enable cross-validation of positive and negative controls in SARS-CoV-2 molecular tests

**DOI:** 10.1101/2020.06.09.143412

**Authors:** Bindu Swapna Madala, Andre L. M. Reis, Ira W. Deveson, William Rawlinson, Tim R. Mercer

## Abstract

DNA synthesis *in vitro* has enabled the rapid production of reference standards. These are used as controls, and allow measurement and improvement of the accuracy and quality of diagnostic tests. Current reference standards typically represent target genetic material, and act only as positive controls to assess test sensitivity. However, negative controls are also required to evaluate test specificity. Using a pair of chimeric A/B RNA standards, this allowed incorporation of positive and negative controls into diagnostic testing for the Severe Acute Respiratory Syndrome Coronavirus-2 (SARS-CoV-2). The chimeric standards constituted target regions for RT-PCR primer/probe sets that are joined in tandem across two separate synthetic molecules. Accordingly, a target region that is present in standard A provides a positive control, whilst being absent in standard B, thereby providing a negative control. This design enables cross-validation of positive and negative controls between the paired standards in the same reaction, with identical conditions. This enables control and test failures to be distinguished, increasing confidence in the accuracy of results. The chimeric A/B standards were assessed using the US Centers for Disease Control real-time RT-PCR protocol, and showed results congruent with other commercial controls in detecting SARS CoV-2 in patient samples. This chimeric reference standard design approach offers extensive flexibility, allowing representation of diverse genetic features and distantly related sequences, even from different organisms.

## INTRODUCTION

Reference standards are required to validate the performance of any diagnostic test [1]. The recent advent of DNA synthesis enables the rapid development of reference standards as synthetic constructs representing target genetic material. These can be used as positive controls to assess sensitivity of the molecular test undergoing evaluation. However, separate negative controls, without the target sequences, are also required to ensure the specificity of the diagnostic test. Reference standards must be validated and proven fit-for-purpose before used in diagnostic tests. In the case of RNA standards, the synthetic controls undergo degradation over time, and can be contaminated, confounding the interpretation of test results. However, this failure of either positive or negative controls is difficult to distinguish from the failure of the diagnostic test itself. For example, if the test returns a negative result from the positive control, it could be because (i) the test failed, (ii) the reference control failed or (iii) a technical issue with the testing platform. This leads to delays in diagnosis, missed diagnoses and invalidation of correct test results.

The use of *in vitro* synthesis of RNA and DNA standards allows flexibility in control design and tailoring of controls to the diagnostic test and targets. Here, we propose a new design strategy for reference standards that uses matched chimeric synthetic standards in accordance with the principle of A/B testing. In this design, all the target sequences of a molecular test are retrieved and split between groups A and B, which are then joined in tandem to form single chimeric sequences A and B. This means that for each target used in the molecular test, standard A would act as positive control, while standard B would act as negative control, or vice-versa. Furthermore, the equally partitioning of target sites between standards A and B enables cross-validation of positive and negative controls, increasing the confidence in test results. Among the benefits of this design, a chimera allows concurrent testing of disparate target regions of a single pathogen or even different organisms and splitting targets between standards A and B enables control cross-validation, facilitating the distinction of control failure from test failure or success.

The recent emergence of the SARS-CoV-2 pandemic has required widespread diagnostic testing for active virus infections, including genome sequencing, but predominantly using real-time reverse-transcriptase polymerase chain reaction (real-time RT-PCR)-based assays [2-4]. The World Health Organisation (WHO) published seven diagnostic testing protocols for detection of SARS-CoV-2 that have been rapidly adopted worldwide, with over 20 million molecular tests performed globally by mid-2020 [5, 6]. These tests typically employ multiple primer pairs homologous with SARS-CoV-2 genes E, N, Orf 1a/1b and RdRp [7-9]. Diagnosis is considered positive if all targets are amplified or presumptive positive if some but not all targets are detected. In addition, some tests contain primer pairs also targeting human genes as internal positive controls to ensure sample quality.

We used a pair of chimeric A/B standards for the WHO-endorsed real-time RT-PCR tests to demonstrate the utility of the chimeric A/B approach to designing reference test standards. Each standard included regions of the coronavirus genome (SARS-CoV-2) that are targeted by published primer pairs. As a result, it is compatible with endorsed diagnostic tests licensed globally. We compared the performance of the synthetic controls to other reference materials and patient samples, and demonstrated how the two synthetic RNA standards can be used to validate the standard real-time RT-PCR test (CDC), and also considered the utility of these standards in other assays [10].

## RESULTS

### Design of RNA standards

DNA synthesis enables rapid and flexible assembly of reference standards, including sequences not present in natural organisms. This allows sequences from different genome regions to be aggregated to address specific requirements in a diagnostic assay. To demonstrate this approach, we designed synthetic reference sequences that encompass the primer binding sites of all WHO-published real-time RT-PCR tests.

We first retrieved the SARS-CoV-2 genome sequence (isolate Wuhan-Hu-1, NC_045512.2), as well as the primer sequences published by the World Health Organisation (WHO) for China, Hong Kong, Thailand, United States (CDC), Germany and France (**Fig. 1a**). Each available real-time RT-PCR test typically comprises 2-3 primer pairs that target different regions of the SARS-CoV-2 genome (**see Supplementary Table 1**). We then aligned the primer pairs to the SARS-CoV-2 genome and identified the coordinates of the amplicons, which were then retrieved along with an additional 30 nucleotides (nt) on either flanking side (**Fig. 1a**).

**Figure 1.**
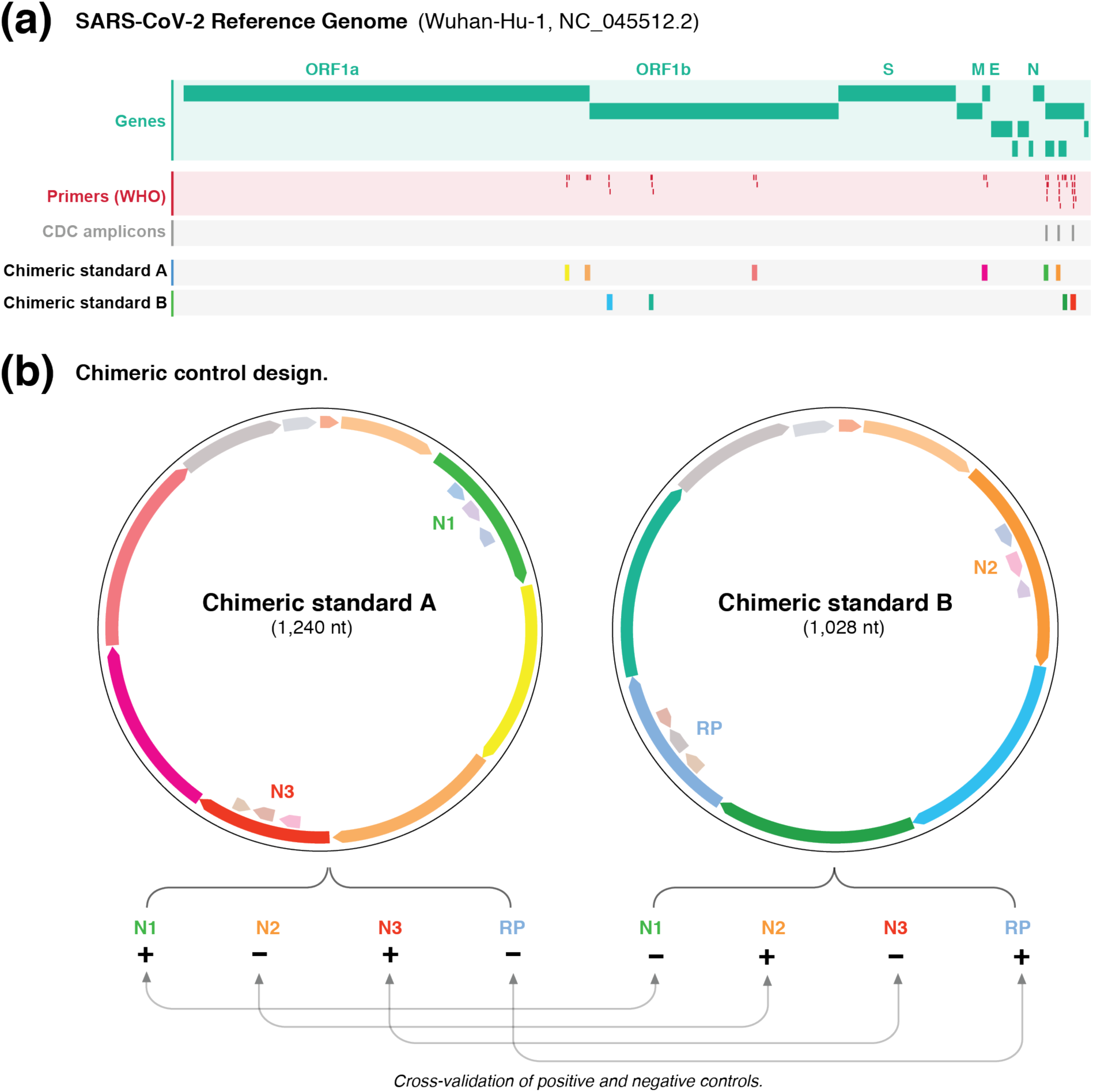
Design of chimeric controls for SARS-CoV-2. **(a)** Genome browser view of the SARS-CoV-2 genome (green) showing WHO-published real-time RT-PCR primer binding sites (red). The expected amplicons for the CDC test are shown in darker grey. The other targeted regions were exclusively partitioned between chimeric A/B standards. **(b)** The different targeted regions for standards A and B were shuffled and joined together to form chimeric sequences. The paired design of chimeric A/B standards, where a target in A is absent in B (and vice versa), enables the synthetic RNA transcripts to simultaneously act as positive and negative controls for the real-time RT-PCR primer/probe sets. This enables internal cross validation of positive and negative controls between standards A and B. The vector backbone was omitted from the representation of the chimeric A/B standards.

We next organised these sequences across two different controls (termed chimeric A/B standards). We partitioned the different targeted regions used by each country into two independent groups and then assembled the regions in tandem (**Fig. 1b**). A fragment of the human RNase P gene (RP), which is used as a positive human control, was also added to the chimeric standard B.

An additional unique control sequence (UCS) was also included at the 5’ region of each standard to enable the unique detection of the standards if required. Each standard sequence was then preceded by a T7 promoter to enable *in vitro* transcription, and followed by a poly-A tract (30nt length) and a restriction site (EcoR1) to enable vector linearization.

The two distinct chimeric A/B standard sequences were then synthesised and cloned into pMK vector backbones (**see Methods**). We then linearized the plasmids and performed *in vitro* transcription to produce the synthetic RNA standards (**Fig. S1**). The resulting RNA transcript products were then purified and validated (see **Methods** and **Supplementary Methods** for detailed protocol).

### Validation of chimeric RNA standards to alternative reference controls

We next validated the chimeric A/B standards by comparison to alternative reference controls. We first performed real-time RT-PCR test using the established CDC primers and protocol [10]. Specifically, this employs CDC primers (IDT) *N1, N2* and *N3* targeting the N gene from SARS-CoV-2 and the human *RNase P* gene. Chimeric standard A includes regions of the *N1* and *N3* targets, while chimeric standard B includes regions of the *N2* and *RP* targets. This allows for cross-validation between the chimeric A/B standards, since the standards alternatively act as positive and negative controls to each primer/probe set in the real-time RT-PCR test.

The real-time RT-PCR was initially performed on the chimeric controls alone. We prepared 10-fold dilutions for each control, starting at 3.96 × 10^8^ copies/μl for A and 4.22 × 10^8^ copies/μl for B (see **Methods**). As anticipated, in the reactions containing standard A, *N1* and *N3* primers returned positive results, while N2 and RP were undetected (**Fig. 2a**). In contrast, in reactions containing standard B, *N2* and *RP* primers returned positive results, while *N1* and *N3* were undetected (**Fig. 2a**). In the real-time RT-PCR reactions with positive results, there is an average increase in Ct values of 3.47 (sd= 0.34) for a 10-fold dilution. These results show that the chimeric controls enable positive and negative cross-validation of the published CDC real-time RT-PCR test for SARS-CoV-2.

**Figure 2.**
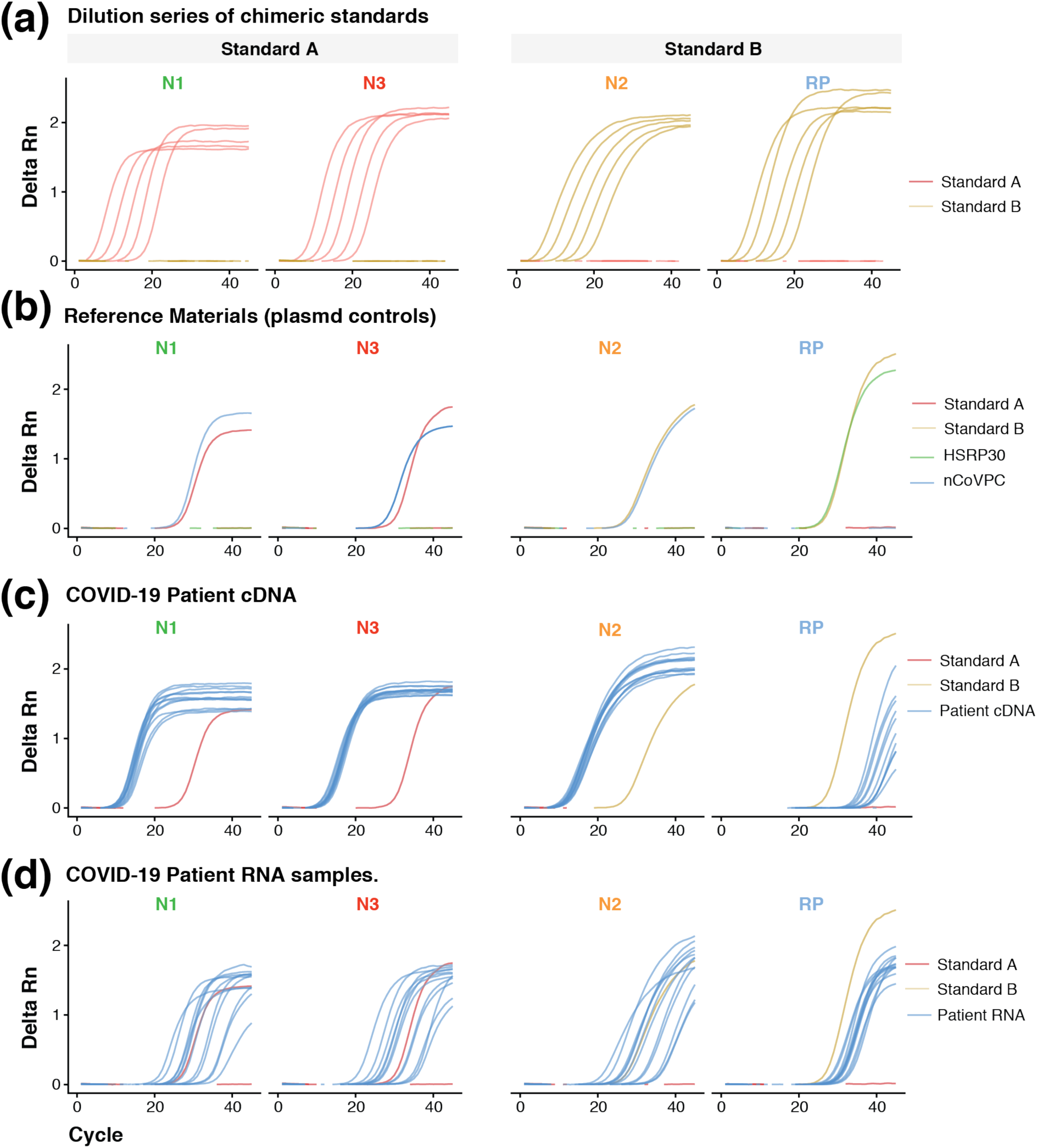
Real-time RT-PCR validation of chimeric A/B standards. Amplification curves for the target genes (N1, N2, N3 and RP) in the CDC real-time RT-PCR test for SARS-CoV-2. (**a**) The results for the chimeric standards alone show that N1 and N3 were detected in standard A, but absent in B, while N2 and RP were detected in standard B, but absent in A. (**b**) The chimeric standards achieve similar results to IDT controls, which provide separate positive (nCovPC) and negative controls (HSRP30). (**c**) Chimeric standards compared to COVID-19 patient samples, where SARS-CoV-2 genome was previously amplified. (**d**) Chimeric standards compared to RNA from confirmed COVID-19 RNA patient samples.

We next performed the real-time RT-PCR test including commercial controls (Integrated DNA Technologies Ltd.) for comparison with the chimeric A/B standards. The positive and negative commercial controls are provided as separate plasmids. The positive control (2019-nCoV_N_Positive Control) contains the complete nucleocapsid gene for SARS-CoV-2, while the negative control (Hs_RPP30) contains a fragment of the human RNase P gene.

Therefore, IDT positive control, 2019-nCoV_N_Positive Control, should detect *N1, N2* and *N3*, but not *RP*, while, IDT negative control, *Hs_RPP30*, should only detect RP. We diluted each plasmid to 4,000 copies/ul along with the chimeric A/B standards and performed the real-time RT-PCR test with the CDC primers and probes. The IDT controls worked as expected and the chimeric A/B standards achieved similar results (**Fig. 2b**). For each of the target genes, The Ct values were comparable between the IDT controls (N1=26; N2=28; N3=28; RP=27) and the chimeric A/B standards (N1=27; N2= 27; N3= 30; RP=27).

### Comparison of chimeric RNA standards COVID-19 patient samples

To validate the commutability of the chimeric A/B standards, we compared their performance in amplifying genomes from 12 COVID patient samples that had been independently diagnosed and sequenced (**see Methods**). These samples were available as raw total RNA containing the viral genome and cDNA, where the viral genome was previously amplified. Therefore, the real-time RT-PCR test with CDC primers should yield positive results for all target genes (*N1, N2, N3* and *RP*) in patient RNA samples and primarily positive results for *N1, N2* and *N3* in patient cDNA samples.

We diluted patient cDNA samples so that on average reactions had 2.79 ng (sd= 0.35) of input DNA. We did not measure the concentration of patient RNA samples due to insufficient starting materials. The real-time RT-PCR amplification results showed that both chimeric A/B standards and patient samples were diagnosed as expected. Target genes *N1, N2* and *N3* were amplified in both patient cDNA (avg Ct; N1=12 ± 0.72, N2=12 ± 0.70, N3=13 ± 0.69) and patient RNA (avg Ct; N1=29 ± 4.85, N2=29 ± 4.92, N3=29 ± 4.89) samples, with the former achieving significantly lower and less variable Ct values (**Figs. 2c**,**d**). However, for the *RP* target gene, which is a positive control for human samples, the Ct value for patient cDNA was significantly higher, since those samples are depleted of human material (37 ±1.83 and 31 ± 1.54, respectively).

To determine the limit of detection (LoD) for the chimeric A/B standards, we performed 100-fold serial dilutions with three technical replicates. As a baseline, we spiked the standards A (3,964 copies/μl) and B (4,221 copies/μl) into separate background samples consisting of the human universal RNA (100 ng). We then performed real-time RT-PCR using the CDC protocol (see **Methods**). As a result, we detected both standards A and B until 10^−4^ dilution, which corresponds to approximate LoD of 0.39 and 0.42 copies/μl, respectively (**Fig. 3a**). As a positive control, the RP primer targeting the human *RNase P* gene were successfully detected in all tested dilutions, for both standards A and B (**Fig. 3b**). Interestingly, N1 primers appear to be more efficient than N3 primers in estimating the LoD for standard A, since the Ct values for N1 (10^0^=21.75 ± 0.09, 10^−2^= 28.26 ± 0.04 and 10^−4^= 34.33 ± 0.96) are significantly lower than N3 (10^0^=25.04 ± 0.06, 10^−2^= 31.33 ± 0.04 and 10^−4^= 37.16), in every dilution, across replicates (**Fig. 3b**).

**Figure 3.**
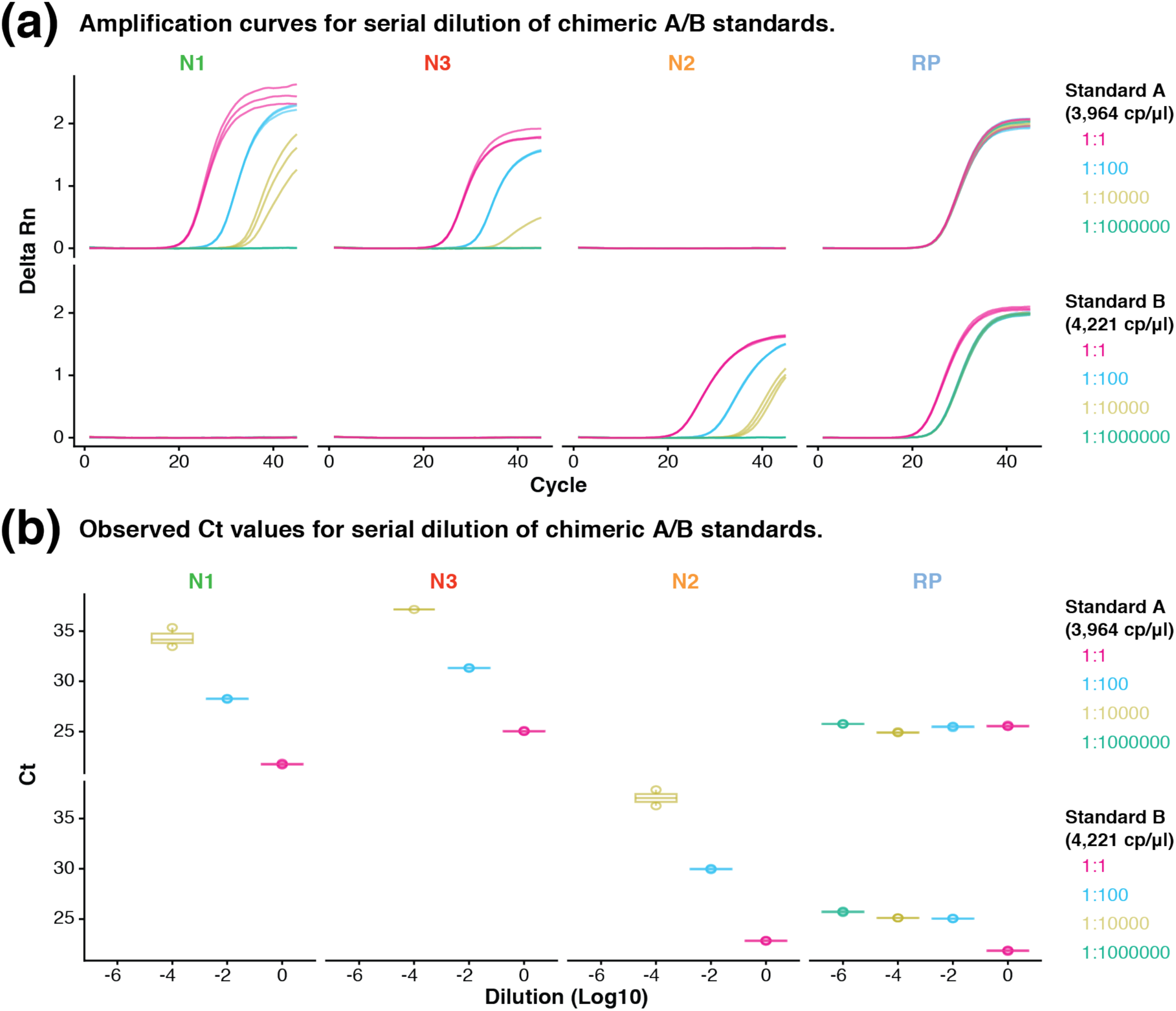
Limit of detection of chimeric A/B standards. (**a**) Amplification curves of targets *N1, N2, N3* and *RP* in 100-fold serial dilutions of standards A and B against the human universal RNA, with three technical replicates. (**b**) Observed Ct values for targets N1, N2, N3 and RP at different dilutions of standards A and B (10^0^, 10^−2^, 10^−4^ and 10^− 6^), across three technical replicates.

## DISCUSSION

The advent of routine DNA synthesis has enabled rapid provision of synthetic reference standards that can be used to validate the accuracy of diagnostic tests. The synthesis of DNA provides a flexible platform to manufacture different reference standards, including non-natural designs. In this case, a single standard can be designed to contain distant genomic regions or even sequences from different organisms. This allows multiple sequences of interest to be included and organised within a single chimeric standard according to the specific requirements of a diagnostic assay. In this study, we use this approach to generate chimeric RNA transcripts that encode different SARS-CoV-2 genomic regions targeted by WHO sanctioned molecular diagnostic tests.

Furthermore, we show how chimeric standards enables matched A/B testing of each primer/probe set used in the real-time RT-PCR. Because the target regions are exclusively distributed between the A/B standards, each inevitably functions as an independent positive or negative control for a given primer/probe set. Additionally, the balanced distribution of target regions for each WHO sanctioned test between A/B standards, ensures cross-validation of positive and negative controls. This improves the confidence in the results of the controls and prevents a test failure being confused with a control failure.

Within this study we successfully validated the chimeric A/B standards using the CDC real-time RT-PCR test for SARS-CoV-2 detection [10]. However, the chimeric A/B standards also contain the target sites in accordance to official guidelines for molecular testing in China, Thailand, Hong Kong, Germany and France, besides those of the United States. This means that the chimeric A/B standards can not only be used as test controls in each of those countries individually, but also provide a common reference to compare test results and the efficiency of their different primer/probe sets.

The Covid-19 pandemic has led to rapid development of novel diagnostic technologies. Design flexibility in DNA synthesis enables the development of reference standards that simultaneously and specifically address the validation requirements of a diverse range of diagnostic tests. We demonstrate the concept with the chimeric A/B standards, but we anticipate this design approach can be used in applications beyond SARS-CoV-2 diagnostic tests. For instance, a single synthetic control could represent multiple genetic features, such as mutations, different haplotype blocks or viral sequences. This can also include microbial sequences, cancer mutations and expressed gene signatures. In future testing of populations with decreased incidence and prevalence of SARS CoV-2 infection and COVID-19 disease, internal controls will be critical in allowing accurate and rapid assessment of infection. This is because the predictive value of molecular and other testing is lower in low prevalence and low incidence populations. In summary, the chimeric A/B approach provides a new model for the development of cost-effective reference standards that allow for controlling of multiple experimental variables based on simple, but comprehensive designs.

## ACKNOWLEDGEMENTS

We thank our colleague Prof Bill Rawlinson for providing patient isolates collected by his laboratory for a separate study. We acknowledge the following funding sources: National Health and Medical Research Council (NHMRC grants APP1108254, APP1114016, APP1136067), UNSW Tuition Fee Scholarship (TFS; to A.L.M.R) and Cancer Institute NSW Early Career Fellowship 2018/ECF013 (to I.W.D.). The contents of the published materials are solely the responsibility of the administering institution, a participating institution or individual authors, and they do not reflect the views of the NHMRC or CINSW.

## Author Contributions

B.S.M, A.L.M.R. & T.R.M. conceived the project and devised the experiments. B.S.M conducted laboratory experiments. I.W.D. and W.R. provided COVID-19 patient samples. A.L.M.R & T.R.M. performed data analysis. A.L.M.R & T.R.M. prepared the manuscript, with support from all co-authors.

## Ethics Declarations

All procedures performed with human participants were in accordance with the ethical standards of the Human Research Ethics Committee at South Eastern Sydney Local Health District (SESLHD), under approval 2020/ETH00287. Where relevant, informed consent was obtained from individual research participants.

## MATERIALS AND METHODS

### Covid-19 Patient samples

Patient samples used for validation tests were collected at the SAViD laboratories at Randwick as part of a quality assurance study. The samples constitute viral RNA extracts (Roche MagNA Pure extraction kit) on nasopharyngeal swabs from patients testing positive for SARS-CoV-2 infection. cDNA was generated from the same isolates using Thermo Fisher Superscript IV VILO Master Mix, according to the recommended protocol. cDNA was amplified with each of 14 x ∼2.5 kb amplicons tiling the SARS-CoV-2 genome, according to a custom protocol [11]. Amplicons were then cleaned with AMPure beads and pooled at equal abundance.

### Commercial controls

We acquired control plasmids from IDT Technologies to be used in the CDC real-time RT-PCR diagnostic assay of SARS-CoV-2. The positive control (2019-nCoV_N_Positive Control) contained the complete nucleocapsid gene, while the negative control contains a portion of the human RPP30 gene. The stocks for each of the plasmids were delivered at 200,000 copies/μl in IDTE pH 8.0. For the real-time RT-PCR, we diluted the plasmids to 4,000 copies/μl each.

### Synthesis and preparation of chimeric standards

The A/B standards were synthesized by a commercial vendor (ThermoFisher – GeneArt) and cloned into pMK vectors. The plasmids containing the standards were each resuspended in 50 μl nuclease free water and transformed in *E. coli* as per manufacturer’s protocol (α-Select Competent Cells, Bioline, Australia). The transformed cells were grown overnight (37°) in LB agar plate containing Kanamycin (100 μg/ml), after which colonies were selected and further cultured overnight (37°; 200 rpm) in 3 ml LB broth also containing Kanamycin (100 μg/ml). The plasmids with the A/B standards were then extracted and purified using the ZymoPURE Plasmid Miniprep Kit (Zymo Research), according to the manufacturer’s protocol. Purified plasmids were linearized by overnight digestion (37°) with EcoRI-HF (NEB) and the products were then visualized on 1% agarose gel. The linear A/B standards were finally treated with Proteinase K and further purified with the Zymo ChIP DCC-25 purification kit (Zymo Research). The final A/B standards were quantified using Qubit dsDNA HS Assay on Qubit 2.0 Fluorometer (Life Technologies) and verified on the Agilent TapeStation with the High Sensitivity DNA Screen Tape Analysis (Agilent Technologies).

### *In vitro* transcription

The ChIP purified A/B standards were submitted to an *in vitro* transcription reaction, incubated overnight at 37°, using the MEGAscript T7 Transcription kit (ThermoFisher) according to the manufacturer’s protocol. The resulting product was then treated with Turbo DNase and the remaining RNA was purified with the Zymo RCC-25 column purification-25 kit (Zymo Research). The A/B RNA standards were quantified using Qubit RNA HS Assay on Qubit 2.0 Fluorometer (Life Technologies) and then verified on the Agilent TapeStation (Agilent Technologies) with the RNA ScreenTape Analysis (Agilent Technologies).

### Quantitative real time PCR

Twenty μl reactions were prepared containing 5 μl of input RNA (patient samples, A/B standards or IDT controls), 5 ul of TaqPath™1-Step RT-qPCR Master Mix, 1.5 μl of the combined CDC primers/probe set and 8.5 μl of Nuclease-free water. Thermo cycling was performed at 25° for 2 min to allow UNG incubation, followed by 15 min at 50° for reverse transcription, then 2 min at 95° for enzyme activation and finally 45 amplification cycles at 95° for 3 seconds and 55° for 30 seconds. The experiment was performed on QuantStudio 7 Flex real-time PCR systems (Thermo Fisher).

### Dilution series

We performed two different serial dilution experiments with the chimeric A/B standards. The first was a 10-fold serial dilution of standards A and B alone, to test their performance in the real-time RT-PCR assay. The baseline concentration was 3.96 × 10^8^ copies/μl for standard A and 4.22 × 10^8^ copies/μl for standard B. We diluted each standard until 10^−5^. The second serial dilution was to estimate the limit of detection (LoD) for the chimeric A/B standards. For this experiment, standards A and B were individually spiked into universal human RNA samples. The baseline concentration was 3,964 copies/μl for standard A and 4,221 copies/μl for standard B and they were each added into 100 ng of universal human RNA. We made 100-fold dilutions for the A/B standards until 10^−6^, and the experiment was performed in three technical replicates.

