## Supplement Materials for "Chimeric synthetic reference standards enable cross-validation of positive and negative controls in SARS-CoV-2 molecular tests"

### TABLE OF CONTENTS

**Supplementary Figure 1.** **P2**

### APPENDIX

**Supplementary Table 1 -WHO endorsed real-time RT-PCR prime/probe sets per country.**

**(a)** **Isolation of chimeric standards A and B plasmids and digestion with EcoRI (different exposures).**

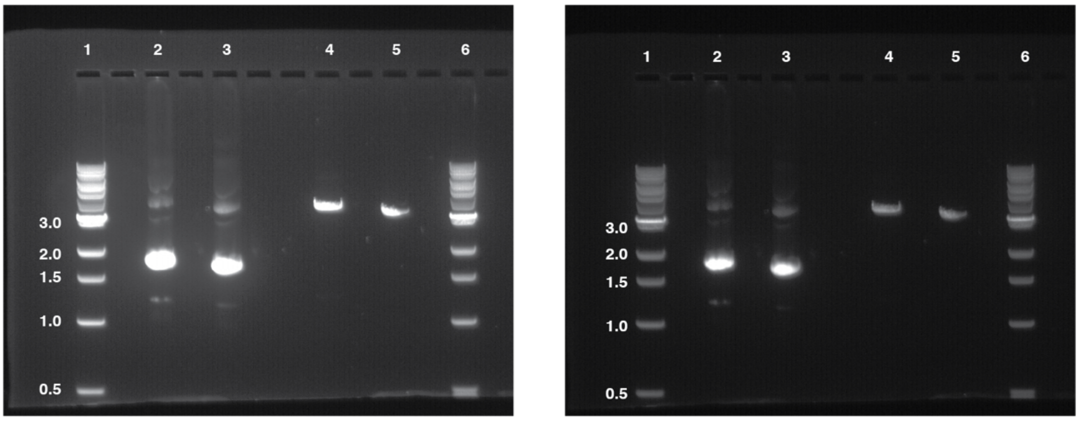


**1- 1Kb ladder; 2- Standard A miniprep; 3- Standard B miniprep; 4- Standard A digestion; 5- Standard B digestion; 6- 1Kb ladder**

**(b)** **TapeStation quantification of *in vitro* transcribed RNA for chimeric standard A.**

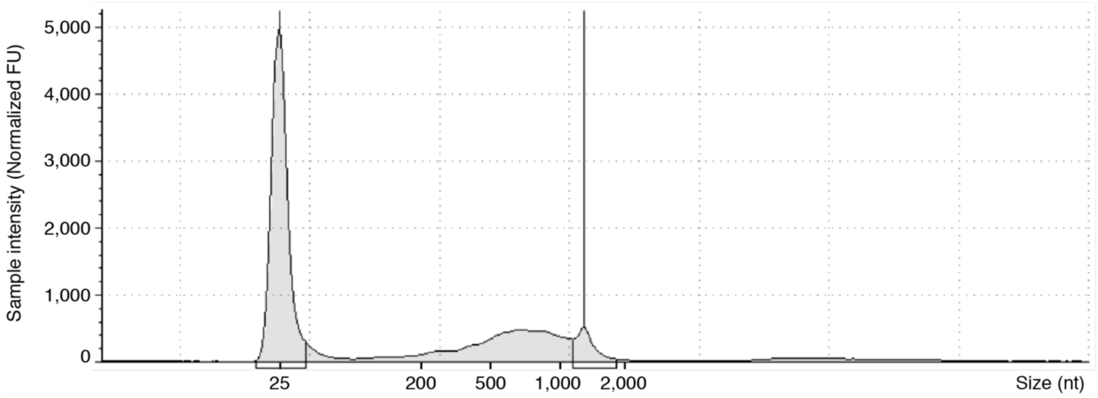


**(c)** **TapeStation quantification of *in vitro* transcribed RNA for chimeric standard B.**

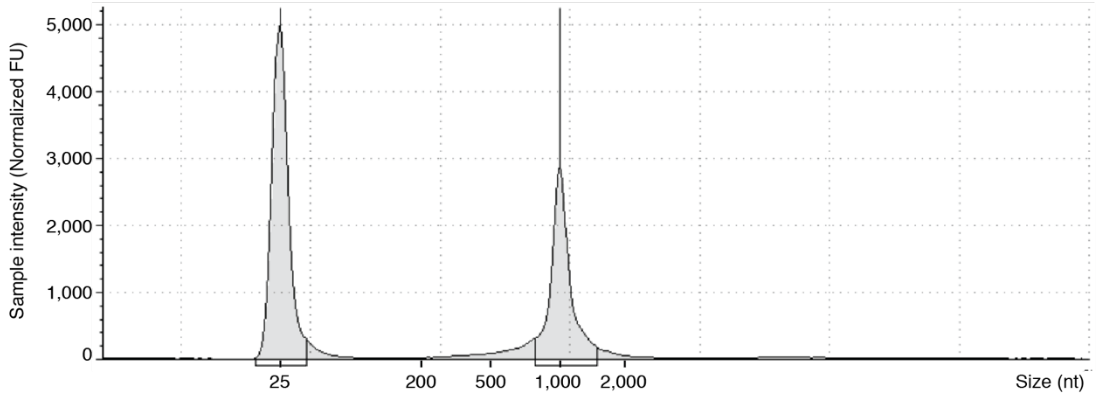


**Supplementary Figure 1**. (**a**) Gel electrophoresis showing miniprep results for chimeric A/B standards and digestion with EcoRI restriction enzyme. Those are photos of the same gel at different exposures. (**b**) TapeStation results for the purified RNA product, resulting from the *in vitro* transcription of the linearized standard A. (**c**) TapeStation results for the purified RNA product, resulting from the *in vitro* transcription of the linearized standard B.
